# Prevalent glutamyl-endopeptidases in the commensal skin microbiome have itch-relevant activity

**DOI:** 10.64898/2026.07.17.739094

**Authors:** Jan-Philipp Wittlinger, Sophie Weninger, Joana Séneca, Aldine Tu, Lukas Grey, Stefan Heber, Michael Fischer, Sven Schneider, Julia Eckl-Dorna, Georg Stary, Thomas Böttcher, David Berry

## Abstract

Atopic dermatitis (AD) is frequently accompanied by pruritus, which has predominantly been attributed to skin colonization by *Staphylococcus aureus*, particularly through cleavage of protease-activated receptor 1 (PAR1) by the glutamyl endopeptidase (GEP) V8 protease. Whether related GEPs from other skin-associated staphylococci contribute to this process remains unclear. Here, we analyzed 273 staphylococcal isolates from skin swabs of 10 AD patients with pruritus, dominated by *S. aureus* and *Staphylococcus epidermidis*. Genome mining using a custom hidden Markov model identified 678 candidate GEPs, which were clustered and resolved into five structurally distinct protease architectures. Representative proteases (V8, Esp, SplB, Csp, and Hsp) were expressed and characterized. Esp displayed GEP activity and PAR1 tethered-ligand cleavage comparable to V8, generating noncanonical cleavage products, while Csp cleaved PAR1 with reduced but substantial efficiency. All representative proteases significantly disrupted barrier integrity in an epithelial barrier model. Analysis of isolate genomes and publicly available *Staphylococcus* genomes showed that V8 and Esp are highly conserved and largely species-restricted, whereas Csp is more broadly distributed across species. These findings identify Esp and Csp as functional GEP virulence factors in *S. epidermidis* and *S. capitis*, capable of activating itch signaling and compromising barrier function. Our study suggests that GEP-mediated pruritus and barrier dysfunction in AD may arise not only from pathogens like *S. aureus*, but also from commensals or opportunistic pathogens such as *S. epidermidis* and *S. capitis*.

**Importance:** *Staphylococcus aureus* colonization on the skin is closely associated with itch in atopic dermatitis (AD) through a secreted protease that cleaves PAR1 on sensory neurons. However, AD patients can experience pruritus without *S. aureus* colonization. This raises the question of whether other skin-colonizing staphylococci contribute to this process. We identify glutamyl endopeptidase homologs, including Esp from *Staphylococcus epidermidis* and Csp from *Staphylococcus capitis*, that cleave the same itch receptor and disrupt epithelial barrier integrity. These serine proteases are distributed across staphylococcal species that colonize human skin. Therefore, itch and barrier dysfunction in AD may not be restricted to *S. aureus* but instead arise from the proteolytic activity of multiple staphylococcal species. This functional redundancy means that other staphylococcal species can sustain GEP-driven itch and barrier dysfunction even in the absence of *S. aureus*, suggesting that therapeutic strategies targeting only S. aureus may overlook these alternative drivers of disease.

## 2 Introduction

Atopic dermatitis (AD) is one of the most prevalent chronic inflammatory skin diseases, affecting up to 20% of children and 10% of adults worldwide (1, 2). The condition is defined by recurrent episodes of intense itch, skin barrier dysfunction, and type 2 immune dysregulation, and is driven by an interplay between genetic predisposition, environmental factors, and the cutaneous microbiome (3, 4). Advances in culture-independent sequencing have revealed that the composition of the skin microbiome in AD is distinct from healthy individuals, with a marked reduction in microbial diversity and a shift toward overgrowth of *Staphylococcus* species (5–7).

Among staphylococcal species, *Staphylococcus aureus* has received the most attention. It colonizes the skin of 70% of AD patients during flares, compared to less than 20% of healthy individuals, and its abundance correlates with disease severity (8–10). *S. aureus* can exacerbate disease through the secretion of virulence factors, including toxins, proteases, and superantigens that can amplify cutaneous inflammation and disrupt the epithelial barrier (11–14). Alongside *S. aureus*, commensal coagulase-negative staphylococci (CoNS), including *S. epidermidis, S. hominis,* and *S. haemolyticus*, are also enriched on AD skin, though their role in disease pathogenesis remains less well defined and even sometimes described as beneficial (15–19).

A key mechanism by which *S. aureus* induces itch is via secretion of a glutamyl endopeptidase (GEP) known as the V8 protease (11). This serine protease cleaves protease-activated receptor 1 (PAR1) on sensory neurons, triggering downstream itch signaling independently of the canonical histamine pathway (11). In addition to its neuronal effects, V8 protease degrades structural components of the epithelial barrier, including tight junction proteins, thereby compounding barrier dysfunction (20). While secreted proteases are encoded in a broad range of skin-associated staphylococci, the prevalence of GEPs and the functional capacity of their encoded proteases to modulate host protease-activated receptor signaling and compromise epithelial barrier function remain largely unknown (21, 22).

Despite the prominence of *S. aureus* in the AD literature, a clinically important subset of patients present with active disease in the absence of detectable *S. aureus* colonization (23). Interestingly, itch severity in these patients is not significantly different from that observed in *S. aureus-*positive AD (23), suggesting that other members of the staphylococci-dominated AD microbiome may also be capable of driving itch.

To address this knowledge gap, we isolated diverse staphylococci from the skin of AD patients, performed whole-genome sequencing of isolates to identify GEPs across multiple staphylococcal species, and functionally characterized the candidate enzymes for protease activity, PAR1 cleavage, and effects on epithelial barrier integrity. Our findings suggest that previously overlooked GEPs from non-*aureus* staphylococci may contribute to itch and barrier disruption in AD.

## 3 Results

### 3.1 Diverse V8 homologs identified in skin staphylococci

To investigate the potential for the skin microbiota to be involved in itch generation in atopic dermatitis (AD), we cultured 349 bacterial isolates from skin swabs collected from 10 AD patients with pruritus. Consistent with previous reports of staphylococcal overgrowth in AD, most isolates belonged to the genus *Staphylococcus* (n=273). Whole-genome sequencing resolved the isolates into 13 species, with *Staphylococcus aureus* and *Staphylococcus epidermidis* dominating in nearly equal proportions across the cohort (Figure 1a; Table S1). We then sought to determine how broadly the capacity for itch-relevant protease activity is distributed across the isolates. The *S. aureus* V8 protease, which is a serine protease with GEP activity, has been shown to induce pruritus through PAR signaling. We therefore screened the 273 genomes and identified a total of 9,946 putative serine proteases. As serine proteases are a general class of proteases, we refined our search by using a custom hidden Markov Model to identify putative GEPs, which resulted in 678 significant hits. GEPs possess a highly conserved catalytic triad (Ser-His-Asp) and a structurally stable substrate-binding pocket that confers specificity for cleavage after glutamic acid residues (24). The model effectively captured the conserved catalytic features of GEPs, making it suitable for identifying close homologs for downstream clustering, structure prediction, and experimental validation. To select representative GEPs for experimental validation, sequences were next clustered at 90% amino acid identity using CD-HIT, yielding 42 clusters (Figure S1a, b). As part of candidate validation, all protease sequences were screened for signal peptides to confirm their potential for extracellular secretion. The structure of each representative protein was then predicted to evaluate whether candidates adopt the canonical fold expected of active GEPs. The valine residue within the charge-compensator region is known to be critical for catalytic activity in the V8 protease, so we used its presence, or the identity of the residue occupying the equivalent structural position, as a key criterion for further grouping (25). This analysis resolved the 42 clusters into 5 distinct structural groups, each representing a different architectural variant of the GEP active site (Figure 1b; Figure S1c-e). One representative protein was selected from each structural group for experimental characterization.

**Figure 1.**
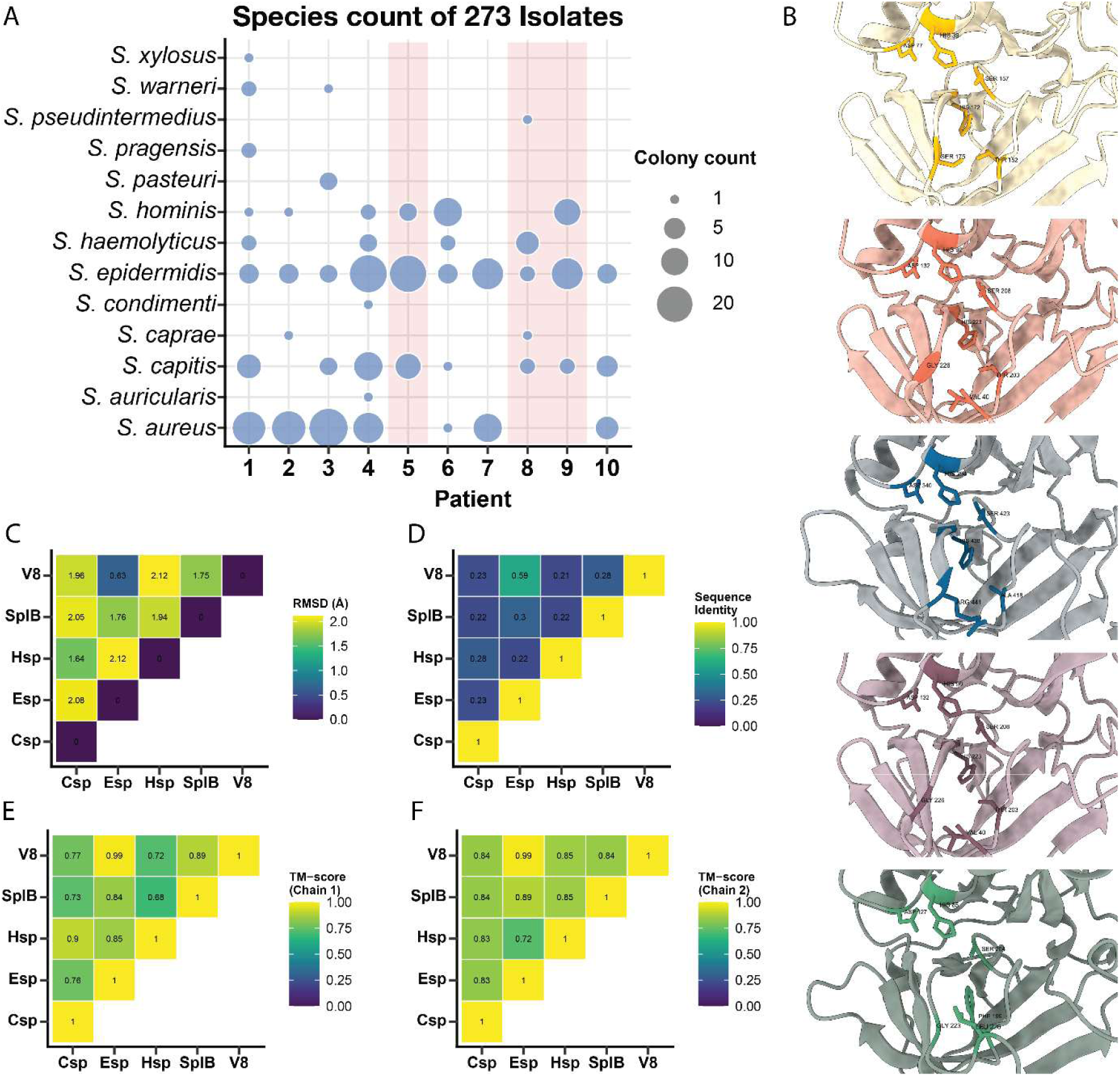
Distribution of staphylococci in atopic dermatitis patients and structural characterization of serine glutamyl endopeptidases. (a) Staphylococcal species counts across 273 isolates from atopic dermatitis patients. Red shading indicates patients from which *S. aureus* was not recovered. (b) AlphaFold2-predicted structures of five serine glutamyl endopeptidases (from top to bottom: SplB, V8, Csp, Esp, Hsp), highlighting the conserved catalytic triad (His, Asp, Ser; sticks) and divergent residues within the substrate-binding cleft. (c-f) Pairwise TM-align-based comparison of the five homologs, showing RMSD (c), sequence identity (d), and TM-scores (e, f) collectively indicating high structural conservation despite low sequence identity.

To further assess the similarity of the five selected proteases, we conducted pairwise comparisons of the homologs (Figure 1c-f). The established candidate V8 and the homolog Esp showed the strongest structural relationship among all pairwise comparisons. Despite a protein sequence identity of 0.59, the structures exhibited a low RMSD of 0.63 Å and exceptionally high TM-scores of 0.99 for both normalizations. This suggests that V8 and Esp fold almost identically and most likely have conserved functional activity. With TM-scores of ∼0.77-0.83 and RMSDs close to 2.0 Å, Csp, on the other hand, had only modest structural similarity to both V8 and Esp. Csp had the highest similarity to Hsp, although comparisons involving Hsp were typically weaker, especially for TM-scores normalized by the larger protein, reflecting that a substantial portion of Hsp’s structure lacks a structural counterpart in the compared protein. Esp-SplB and SplB-V8 had comparatively high TM-scores (0.84-0.89), indicating partial conservation within this subgroup.

### 3.2 Esp from *S. epidermidis* has strong GEP and PAR1 tethered ligand cleavage activities

Supernatants from *S. aureus* (encoding V8 and SplB), *S. epidermidis* (encoding Esp), *S. hominis* (encoding Hsp) and *S. capitis* (encoding Csp) were next analyzed to evaluate secreted GEP activity. Given that the *agr* quorum-sensing systems are present across the tested *Staphylococcus* species and regulate the expression of secreted virulence factors at the late exponential growth phase (26), supernatants were harvested at this growth stage and GEP activity was assessed using a GEP-specific substrate. *S. aureus* and *S. epidermidis* showed strong GEP activity, whereas *S. capitis* and *S. hominis* showed moderate activity (Figure 2a, b). Transcription of the genes of interest was confirmed by RT-PCR (Figure 2c, Figure S2a) and Sanger sequencing of the resulting amplicons.

**Figure 2.**
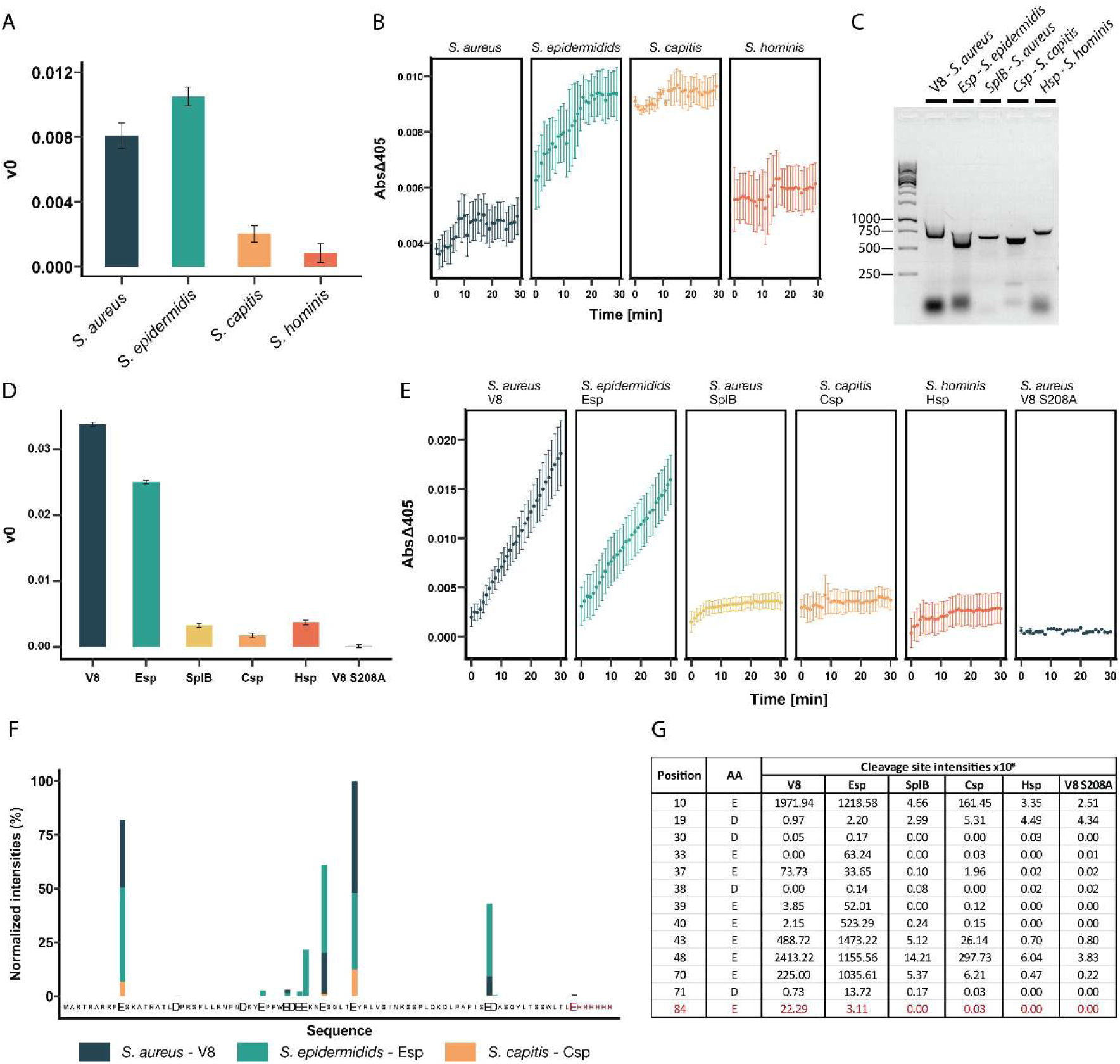
Serine glutamyl endopeptidase expression and activity against synthetic and natural substrates. (a) Initial velocity (v₀) of Z-FLE-pNA substrate cleavage by staphylococcal culture supernatants derived from the linear phase of the reaction. (b) Absorbance at 405 nm over 30 min normalized to LB control containing the chromogenic substrate. (c) Transcriptional confirmation of candidate GEP homologs by RT-PCR using extracted RNA, with their respective size in base pairs confirmed through Sanger Sequencing. (d) Initial velocity (v₀) of Z-FLE-pNA substrate cleavage by recombinant proteins expressed in *E. coli*. (e) Absorbance at 405 nm over 30 min for recombinant proteins normalized to LB control. (f) Cleavage site intensities after the indicated amino acid, normalized to the highest measured intensity. (g) Cleavage-site intensities for proteases at their respective positions. Amino acids and positions colored in red result from the expressed His6-tagged construct and are not native to PAR1.

Having confirmed GEP activity in staphylococcal isolates, we next characterized the activity of the identified proteases. We heterologously expressed the proteases and enzymatically activated them. Non-activated controls underwent identical treatment in the absence of thermolysin. Consistent with computational predictions, Esp displayed a GEP activity profile comparable to V8, supporting its classification as a functional GEP (Figure 2d, e; Figure S2b). The critical role of an intact active site was confirmed by the V8 active-site mutant, which exhibited a complete loss of activity for the synthetic substrate. In contrast, SplB, Csp, and Hsp showed only minimal enzymatic activity.

Given that V8 protease-mediated cleavage of the PAR1 tethered ligand triggers itch signaling, we expressed a codon-optimized construct encompassing the PAR1 tethered ligand domain (PAR1^A22-T102^) to serve as a substrate for detecting protease activity and mapping cleavage sites (11). The expressed construct spanned amino acids A22 through T102, corresponding to positions 1-82 in the substrate map, and was incubated separately with the activated recombinantly produced proteases. The most abundant cleavage sites were identified at E10, E43 and E48. E10 and E48 recorded the highest cleavage intensities for V8 while E43 showed the highest cleavage intensity for Esp (Figure 2f, g; Table S1a). Notably, this hierarchy was consistent across all GEP-specific cleavage events. Whenever cleavage after glutamate was detected, V8 and Esp followed by Csp, generally had the highest cleavage intensities, mirroring the overall activity pattern. In contrast, cleavage at other sites showed more comparable intensities across the homologs, suggesting that these events reflect broader proteolysis rather than specific GEP activity.

### 3.3 V8 protease homologs disrupt epithelial barrier integrity

Having established protease expression and activity for synthetic and natural substrates, we next investigated whether V8 protease homologs contribute to epithelial barrier impairment. PAR1-expressing pruriceptive neurons innervate both the epidermis and dermis, and disruption of the epithelial barrier may therefore be required for access to protease-activated receptors on neurons (11, 27). To validate the assay and determine an effective concentration, we tested a commercially available V8 protease at 10 µM and 2.5 µM alongside a buffer control (Figure S3). Based on these results, 10 µM was selected for subsequent experiments. Monolayers of cultured primary epithelial cells were exposed to proteases, and changes in cell index were monitored using the impedance-based xCELLigence system. All activated proteases significantly reduced epithelial barrier integrity compared to their non-activated counterparts, with no significant pairwise differences in barrier-disrupting activity between any of the activated proteases (two-way ANOVA with Tukey’s posthoc test, *p* < 0.001, Figure 3). These findings demonstrate that GEPs from non-*aureus* staphylococci disrupt epithelial barrier function, indicating a possible role for these proteases in the barrier breakdown observed in atopic dermatitis.

**Figure 3.**
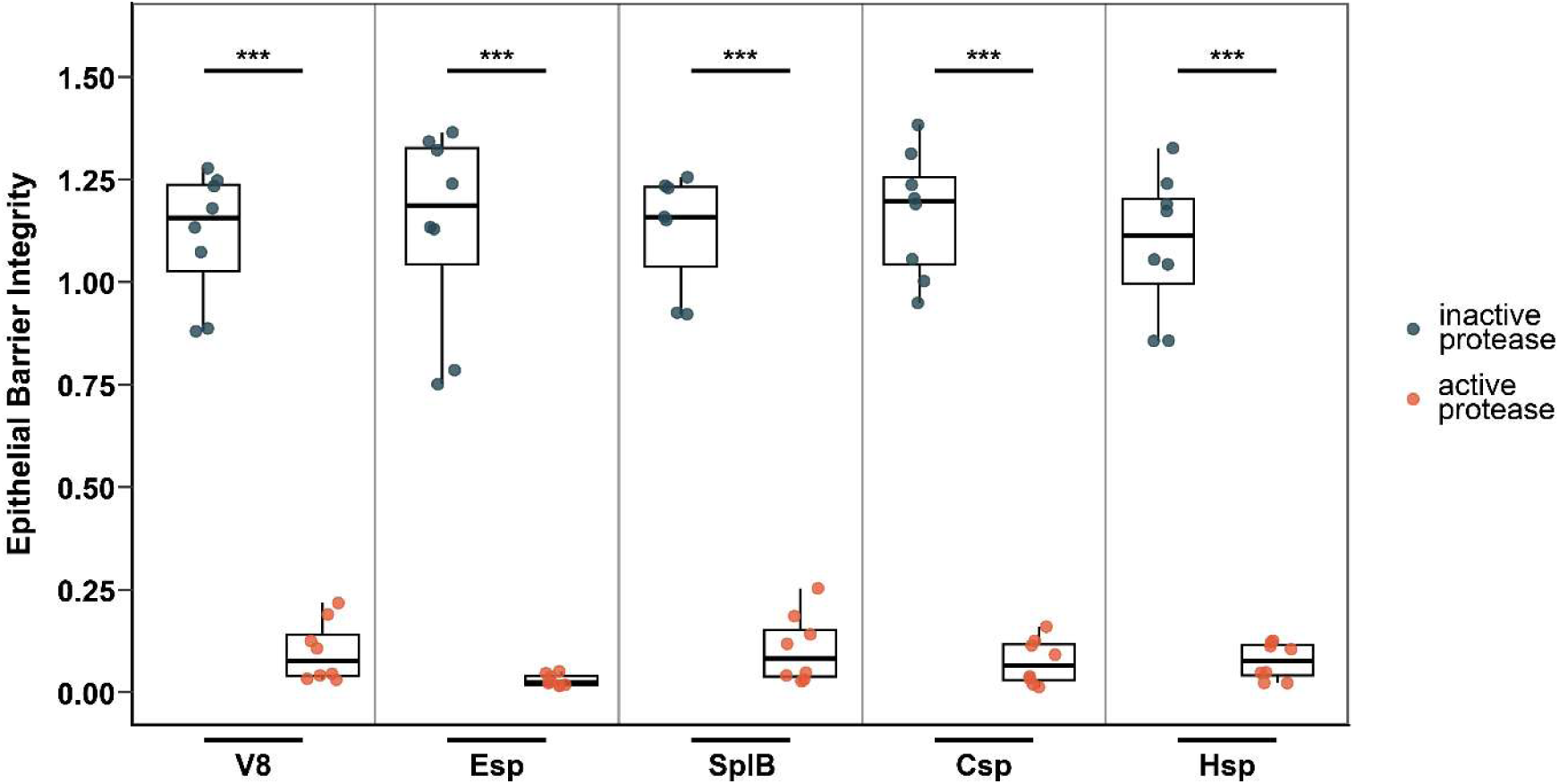
Staphylococcal proteases impair epithelial barrier integrity. Box plots showing the epithelial barrier integrity at the 48-hour time point for cells treated with active or non-activated recombinant proteases (V8, Esp, SplB, Csp, Hsp). Differences between activated protease groups were assessed by two-way ANOVA with Tukey’s *posthoc* test. levels: * *p* < 0.05, ** *p* < 0.01, *** *p* < 0.001. n = 2 technical replicates, n = 4 patients for proteases, except for one patient in SplB (n = 1 technical replicate).

### 3.4 Species-specific distribution and conservation of V8 protease homologs

We next assessed the distribution and conservation of the V8 protease homologs across the genus *Staphylococcus* using our AD patient isolate collection as well as publicly available genomes (Figure 4 a,b). Within our collection, *S. aureus* was the dominant species encoding V8 and the protein was conserved in all but one analyzed genome (Figure 4a). SplB was detected in all *S. aureus* isolates and additionally in a single *S. pseudintermedius* isolate. *S. epidermidis* was the primary species encoding Esp, with one isolate carrying two copies of the gene. Csp was detected frequently in *S. capitis* and *S. haemolyticus* and was also found in *S. condiment* and *S. caprae.* Hsp showed the broadest distribution of all proteins screened, detected across nine species including *S. capitis*, *S. epidermidis*, and at full prevalence in *S. haemolyticus*, *S. hominis, S. condimenti*, *S. pasteuri, S. pragensis, S. warneri*, and *S. auricularis*.

**Figure 4.**
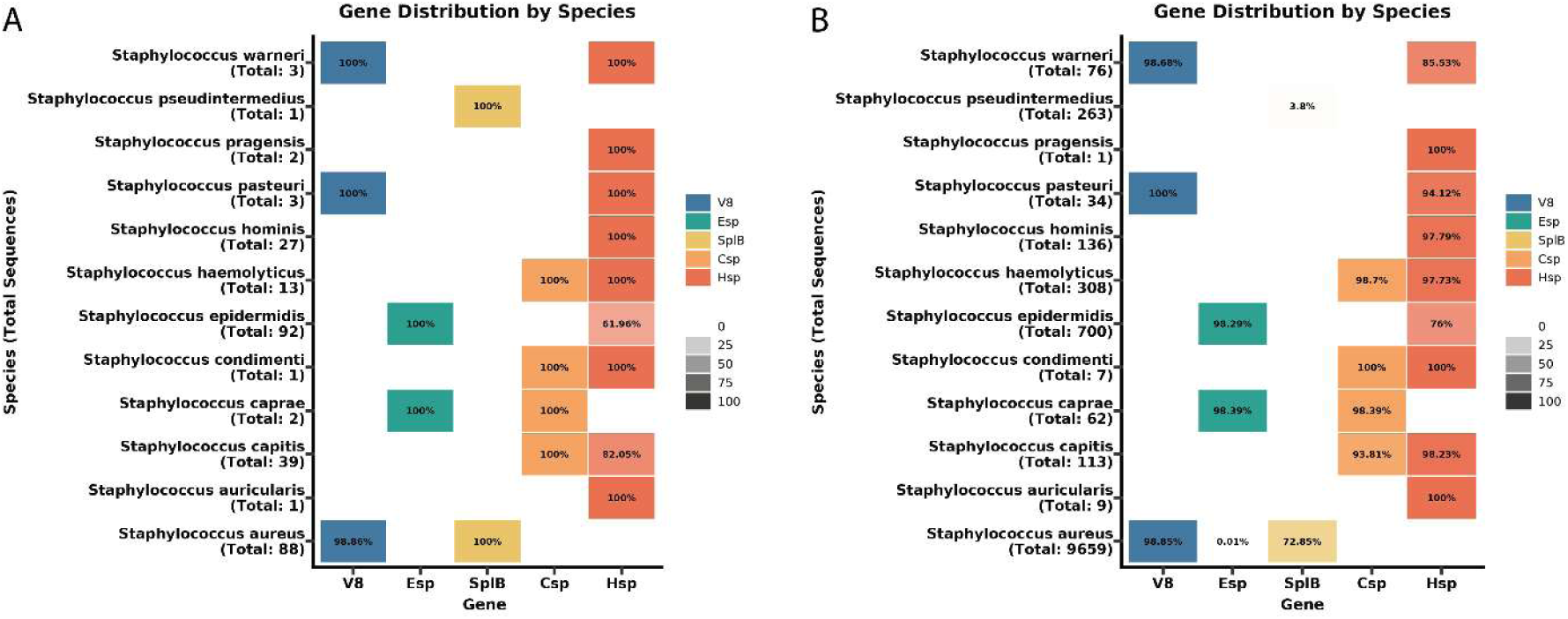
Distribution and conservation of staphylococcal glutamyl endopeptidase homologs across species. Prevalence of V8, Esp Csp, SplB and Hsp proteases across staphylococcal species are shown. Tile shading intensity reflects the proportion of isolates or genomes harboring the respective protease. Empty tiles indicate the absence of a significant hit. (a) Prevalence of proteases across *Staphylococcus* isolates obtained from atopic dermatitis patients. (b) Prevalence of proteases across publicly available *Staphylococcus* genomes (n=12,588), as determined by BLAST screening using V8, Esp Csp, SplB and Hsp as reference protein sequences.

Subsequently, we examined the prevalence and conservation of these proteases in publicly available genomes. A BLAST search was performed using V8, Esp Csp, SplB and Hsp as reference protein sequences and a total of 12,588 *Staphylococcus* genomes. We observed distinct conservation patterns for each protease (Table S3, Figure 4). V8 was identified in 98.9% of *S. aureus* genomes. V8 homologs were also detected in *S. pasteuri* and *S. warneri*, though with reduced sequence identity and coverage (Table S3). SplB was detected in 72.9% *S. aureus* genomes (72.9%) and 45.4% *S. pseudintermedius* genomes. Esp was present in 98.3% *S. epidermidis* genomes and detected in 98.4% of *S. caprae* genomes, though with lower sequence identity and coverage (Table S3). The Csp protease showed a broad species distribution, being detected across *S. capitis, S. caprae, S. haemolyticus*, and *S. condimenti* (Figure 4b). Hsp exhibited the widest phylogenetic distribution of all proteases screened, with hits detected across nine staphylococcal species. Taken together, V8 and Esp are highly conserved in *S. aureus* and *S. epidermidis*, respectively, SplB is prevalent in most *S. aureus* genomes, Hsp displays the broadest phylogenetic distribution across the genus, and Csp shows a more variable distribution confined to a cluster of phylogenetically related species.

## 4 Discussion

Non-*aureus* staphylococci are often neglected as a potential source of pathogenic effects in AD. In this study, we identified and characterized a family of GEP homologs across skin-colonizing staphylococci. We demonstrated that these proteases are biochemically active, can cleave the PAR1 tethered ligand, and disrupt epithelial barrier integrity, implicating GEPs beyond *S. aureus* as potential contributors to AD pathogenesis.

The *S. aureus* V8 protease can induce itch by cleaving PAR1 (11), making GEPs prime candidates for contributing to the pathogenesis of pruritus in AD. Screening of genomes of staphylococcal isolates from AD patients revealed a diverse array of GEPs not only in *S. aureus* but also across multiple coagulase-negative staphylococcal species, including *S. epidermidis, S. capitis,* and *S. hominis*.

We found that Esp has comparable biochemical activity to the canonical V8 protease. Importantly, Esp cleaved the PAR1 tethered ligand, linking its activity directly to an itch-relevant signaling pathway. Notably, cleavage did not occur at the canonical Arg41-Ser42 site but instead generated two newly exposed N-terminal serine residues analogous to the first residue of the native tethered ligand motif SFLLRN. This is consistent with the study establishing the V8-PAR1 connection, which also reported cleavage at alternative rather than canonical sites, and is further supported by evidence that peptides such as TFRGAP and SFNGGP are sufficient to activate PAR1 (11, 28). This suggests that the noncanonical cleavage products generated here may retain agonistic capacity and thereby has the potential contribute to itch signaling. Beyond Esp, Csp also cleaved the PAR1 tethered ligand with reduced activity, which may be explained by the absence of an N-terminal valine residue that is important for stabilizing the substrate within the active site and which is present in both mature V8 and Esp. Although structural predictions for SplB, Csp, and Hsp show structural similarity to V8 and Esp, residues situated within loops could obstruct the active site and potentially hinder catalytic function. Even though Csp showed only modest activity against the synthetic GEP substrate and in culture supernatants, its cleavage of the PAR1 construct at glutamyl sites suggests that substrate context and local sequence preferences within a physiologically relevant target may facilitate cleavage beyond what bulk activity measurements alone would predict.

While some of the homologous proteases tested lack the specific PAR1 cleavage activity demonstrated for V8, Esp and Csp, this does not exclude them from acting as virulence factors in a more complex cellular context, where a broader range of targets are available. Unlike synthetic substrates or isolated receptor constructs, epithelial tissues contain a wide range of protease-sensitive targets, including tight junction proteins, whose degradation has been proposed as a key mechanism underlying protease-driven barrier dysfunction (29, 30). Indeed, staphylococcal proteases, including V8, have been shown to degrade tight junction components and compromise transepithelial resistance (17, 31, 32). Our findings extend this capacity to GEP homologs from coagulase-negative staphylococci species that were not previously implicated in barrier disruption. Together, these results demonstrate that Esp may be an important GEP virulence factor in *S. epidermidis* and that GEP homologs across multiple staphylococcal species may collectively contribute to the compromised skin barrier in AD, potentially creating a self-amplifying cycle in which barrier breakdown facilitates deeper protease penetration, PAR1 activation on sensory neurons, and further itch-scratch-mediated damage.

The distribution analysis of GEP homologs across our AD patient isolate collection and publicly available staphylococcal genomes revealed distinct conservation patterns, suggesting that our isolates are in respect to the GEP homologs they encode, largely representative of their species. Most strikingly, V8 and Esp emerged as highly species-specific proteases, with V8 present in nearly all *S. aureus* genomes and Esp in all *S. epidermidis*. V8 is part of the *agr*-regulated virulence program in *S. aureus* and plays roles in immune evasion, biofilm dispersal, and host tissue degradation (26, 33). Similarly, the conservation of Esp in *S. epidermidis* suggests that Esp also serves a central role in *S. epidermidis* virulence while there are still contradicting reports about regulation through *agr*-quorum-sensing (34, 35). Although *S. epidermidis* has traditionally been considered a skin commensal, studies indicate that its abundance on the skin of individuals with AD increases during disease flares compared with both post-flare periods and healthy controls (7, 36). Additionally, it has been shown that *S. epidermidis* can disrupt the skin barrier by forming biofilms and expressing the cysteine protease EcpA (17, 37, 38). The widespread presence of Esp in *S. epidermidis* suggests that GEP-driven epithelial barrier disruption and PAR1 cleavage could potentially impact AD patients regardless of *S. aureus* colonization.

The biochemical and genomic evidence presented here establishes Esp as a GEP virulence factor in *S. epidermidis* with relevance to AD pathogenesis. Confirming this role in more complex biological systems will be an important next step. Mouse models of AD and skin equivalents would allow the contribution of Esp to barrier disruption and itch to be assessed in a tissue context that reflects the in vivo situation. Similarly, quantifying Esp transcriptional activity directly in clinical skin swabs from AD patients at different disease stages would establish whether Esp expression correlates with disease severity. Finally, given that the GEP homologs characterized here collectively target both the PAR1 itch axis and epithelial barrier integrity, they represent a pathogenic mechanism across multiple staphylococcal species. This broadens the case for developing protease inhibitors that address the full staphylococcal proteolytic load on AD skin, rather than focusing solely on V8 protein. Overall, our results raise the possibility that microbiome-induced pruritus and proteolytic barrier dysfunction on AD skin may be influenced not only by *S. aureus* but rather by the collective GEP activity of multiple staphylococcal species.

## 5 Data Availability

Genomes sequenced in this study have been deposited in the NCBI repository under BioProject accession PRJNA1455545. Individual genome assembly accession numbers are provided in Supplementary Table S4. The sequences of the proteases analyzed in this study have been deposited under accession numbers SAMN60995552 (V8), SAMN60995453 (Esp), SAMN60995552 (SplB), SAMN60995431 (Csp), SAMN60995536 (Hsp).

The mass spectrometry proteomics data have been deposited to the ProteomeXchange Consortium via the PRIDE (39) partner repository with the dataset identifier PXD079627.

## 6 Material and Methods

### 6.1 Staphylococci isolation

Skin swabs from atopic dermatitis patients were taken from a 1 cm² area and resuspended in PBS by wringing against the tube walls. Samples were stored at −80°C in 25% glycerol. After thawing, 100 μL of undiluted, 1:10, and 1:100 diluted suspensions were plated on Tryptic Soy Agar (Sigma-Aldrich) with 5% defibrinated sheep blood (Thermo Fisher Scientific) and Mannitol-Salt agar (MSA) and incubated at 37°C and 30°C for 24 h. MSA was prepared with proteose peptone (10 g/L, Oxoid), beef extract (1 g/L, Sigma-Aldrich), NaCl (75 g/L, Carl Roth), D-mannitol (10 g/L, Sigma-Aldrich), phenol red (0.025 g/L, Sigma-Aldrich), and agar (15 g/L, Sigma-Aldrich) in distilled water, and the pH was adjusted to 7.4 ± 0.2. Morphologically distinct colonies were picked, re-streaked three times on fresh blood agar, and grown overnight in TSB (37°C, 180 rpm). Glycerol stocks (25%) were prepared and stored at −80°C.

Single colonies, after the third re-streak, were picked and suspended in 50 µL nuclease-free water and 2.5 µL of this suspension was used as template per reaction. PCRs (25 µL) contained 17 µL H₂O, 2.5 µL 10× DreamTaq buffer (1× final), 2.5 µL 2 mM dNTP mix (0.20 mM total dNTPs final), 0.125 µL 27F primer (100 µM, 0.25 µM final, AGAGTTTGATYMTGGCTC), 0.125 µL 1492R primer (100 µM 0.25 µM final, GGYTACCTTGTTACGACTT), 0.25 µL DreamTaq DNA polymerase (5 U/µL; 1.25 U per reaction), and 2.5 µL template. Thermocycling was: 95 °C for 4 min; 30 cycles of 95°C for 30 s, 51°C for 30 s, and 72°C for 90 s; followed by 72°C for 10 min and hold at 10°C. Amplification products were analyzed by agarose gel electrophoresis and sent for Sanger sequencing with Microsynth AG. ARB-Silva database (v. 138.1) (40, 41) was used to identify the isolates at the genus level.

### 6.2 Whole genome sequencing

High molecular weight DNA was extracted using the Monarch gDNA purification kit (NEB), with the specific protocol for Gram positive bacteria. Pellets were initially treated with an enzymatic cocktail (innuPREP Bacteria Lysis Booster, Innuscreen GmbH) to facilitate chemical cell wall lysis. Samples were diluted and equimolarly barcoded using the SQK-RBK114.96 (Oxford Nanopore Technologies) with the following protocol modifications: we increased the sample input to 200ng/sample, added 1.5 µl barcode/sample, and added +0.5 µl of rapid adaptor to the barcoded library. The final library was loaded on a R10.4.1 flowcell (FLO-PRO114, Oxford Nanopore Technologies) and sequenced for on a Promethion P2 solo (Oxford Nanopore Technologies) using Minknow (v. 23.11.7, Oxford Nanopore Technologies). Flowcell light shields were used.

Nanopore reads were based-called using Dorado server basecaller (v. 7.8.2) using super accuracy mode. The nanopore reads were length and quality filtered using chopper (v. 0.7) (minlen=500bp and mean read Qscore >15) and subsequently assembled using flye (v. 2.9.3) (42) with “–nano-hq” and polished once with medaka (v. 1.11.3, github.com/nanoporetech/medaka). The model used was “r1041_e82_400bps_sup_v4.2.0”. Contigs <1000bp were removed. Assemblies were QC’d using QUAST (v. 5.2.0) (43, 44), CheckM (v. 1.2.2) (45), and classified using GTDBtk (v. 2.3.2) (46, 47).

### 6.3 GEP homolog detection

To annotate the genomes Prokka (v1.14.6) (48) was used. A profile hidden Markov model (HMM) was constructed using HMMER (v3.4) (49) to screen genomic sequences for glutamyl endopeptidase (GEP) homologs. Five experimentally confirmed GEP sequences served as the training set: three from *S. aureus* (UniProt: P0C1U8, P09332, P09331), one from *S. epidermidis* (UniProt: P0C0Q1), and one from *Streptococcus griseus* (UniProt: Q54211). The input sequences were aligned using Clustal Omega (v. 1.2.4) (50) with default parameters to generate a high-quality multiple sequence alignment. The resulting alignment served as the input for the hmmbuild command, which was executed with default settings to generate the profile HMM. The profile was then applied to annotated protein datasets using hmmsearch with default settings. To reduce redundancy among the identified homologous sequences, clustering was performed using CD-HIT (v. 4.8.1) (51, 52) with default parameters and a sequence identity threshold of 0.9. From each cluster, one representative protein sequence was selected for downstream functional and structural characterization. The representative sequences were subsequently analyzed for the presence of N-terminal signal peptides using SignalP (v. 6.0h-fast-3.10.14) (53). The tool was executed with default settings. Processed sequences were used for structure prediction using AlphaFold (v. 2.3.2) (54). Identification of start codons for selected protein sequences was performed using interproscan (v. 5.75-106.0-11.0.4) (55). Methionine residues located between predicted disordered regions and associated Pfam domain models were used to confirm the proposed start positions of the mature protein sequences.

### 6.4 Public Database comparison

RefSeq genomes for all *Staphylococcus* species (accessed January 16, 2026) were downloaded. BLAST+ (v. 2.17.0) (56) was used to construct a custom database from these RefSeq genomes, which was subsequently queried to assess the presence of reference protein sequences identified through whole-genome sequencing. Blast results were quality filtered with an e-value of 1e^-5^, identity ≥ 40 % and coverage of ≥ 70 %. Duplicates were resolved by selecting the result with the highest coverage and sequence identity while also having the lowest E-value. In cases where no single result was clearly superior across all three criteria, the hit with the lowest E-value was chosen.

### 6.5 General analysis

If not indicated otherwise the analysis was done in R (v. 4.4.2) (57) with the following packages: patchwork (v. 1.3.2), ggprism (v. 1.0.7), ggplot2 (v. 4.0.1), ggbreak (v. 0.1.6), ggridges (v. 0.5.7), stringr (v. 1.5.1), purrr (v. 1.0.2), tidyr (v. 1.3.2), dplyr (v. 1.1.4), writexl (v. 1.5.4), readr (v. 2.1.6), readxl (v. 1.4.5), tibble (v. 3.2.1), lubridate (v. 1.9.4), forcats (v. 1.0.1), tidyverse (v. 2.0.0) (58), Biostrings (v. 2.74.1) (59),writexl (v. 1.5.4), pwr (v. 1.3-0) (60), DescTools (v. 0.99.61.14) (61)

### 6.6 Reverse Transcriptase PCR (RT-PCR)

For DNA extraction, overnight culture of isolates was pelleted for 3 min at 10,000 × g and resuspended in 1 mL of PBS. The mixture was transferred to a bead-beating tube (lysing Matrix B, MP Biomedicals) and cells were lysed using the MP-FastPrep24 5g (MP Biomedicals) with the following specifications: 6 m/s, 40 s, 2 cycles, 300 s rest time. The lysate was centrifuged at 14000 × g for 10 min. DNA extraction continued using the supernatant according to the innuPREP DNA Mini Kit 2.0 (Innuscreen GmbH) with proteolytic lysis and DNA extraction.

For RNA extraction, overnight culture of isolates was diluted 1:100 in LB and grown at 37°C and 180 rpm. At OD_600_ of 0.8 the culture was centrifuged for 3 min at 10000 × g and resuspended in 800 µL of buffer RL. The mixture was transferred to a bead-beating tube (lysing Matrix B, MP Biomedica*ls*) and cells were lysed using the MP-FastPrep24 5g with the following specifications: 6 m/s, 40 s, 2 cycles, 300 rest time. The lysate was centrifuged at 14000 × g for 10 min. RNA extraction continued using RNase-Free DNase I Kit (Norgen) after mixing the supernatant 1:1 with Ethanol. After the RNA extraction, the DNA was depleted using TURBO DNA-free™ Kit (Invitrogen) and results were checked with PCR using 16S region (27F: AGAGTTTGATCMTGGCTCAG; 1492R: GGTTACCTTGTTACGACTT) with the following cycling conditions: initial denature 95°C for 4 mins, and then denaturing at 95°C for 30 s, annealing at 51°C for 30 s, and extension at 72°C for 90 s, followed by a final extension at 72°C for 5 min.

RT-PCR was performed using iTaq Universal One-Step RT-qPCR Kit (Bio-RAD) according to the manufacturer’s protocol. Cycling conditions for the respective genes were: reverse transcriptase reaction 50°C for 10 min, initial denature 95° C for 1 min, and then denaturing at 95°C for 20 s, annealing at temperature specific for the gene (Table S4) for 30 s, and extension at 72°C for 60 s, followed by a final Extension at 72°C for 5 min.

### 6.7 Expression of GEPs

Plasmid DNA (GenScript) of each protein construct was dissolved in sterile water and transformed into competent *E. coli* T7 Express LysY (NEB) according to the manufacturer’s protocol. In brief, cells were heat-shocked (42°C, 10 s), followed by recovery in LB medium (37°C, 1 h). Transformants were selected on LB agar plates containing ampicillin (100 μg/mL) overnight at 37°C. Single colonies were used to inoculate overnight cultures in ampicillin-supplemented LB medium, and glycerol stocks (25%) were prepared and stored at −80°C.

Overnight cultures of *E*. *coli* T7 Express LysY carrying recombinant plasmids were used to inoculate 500 mL ampicillin-supplemented (100 μg/mL) LB medium. Cultures were grown at 37°C to OD_600_ = 0.3, induced with 0.2 mM IPTG, and incubated overnight at 18°C. Cells were harvested by centrifugation (4500 rpm, 20 min, 4°C), washed with 1x PBS, and stored at −80°C. Cell pellets were resuspended in 1x PBS and lysed by sonication (Branson Digital Sonifier; 10% amplitude, 3x 10 pulses, 0.5 s on/1.5 s off). Lysates were clarified by centrifugation (full speed, 30 min, 4°C), sterile filtered, and kept on ice until purification by affinity chromatography.

His-tagged proteins were purified by immobilized metal affinity chromatography using a 5 mL HiTrap TALON crude column on an ÄKTA start system (Cytiva) with all Buffers used according to the manufacturers protocol. Lysates were applied in binding buffer, washed with 10 column volumes of washing buffer, and eluted in 6 column volumes of elution buffer, collecting 1 mL fractions. Fractions were analyzed by SDS-PAGE (55 mA, 300 V) using InstantBlue® Coomassie staining (Abcam) and an Unstained Protein MW Marker (Thermo Fisher). Fractions containing the target protein were pooled and concentrated using 3 kDa centrifugal filters (Amicon Ultra, 5000 × g). Buffer exchange into 1× PBS was performed using PD-10 desalting columns (Sephadex G-25, Cytiva). Protein concentration was determined using a Pierce BCA Protein Assay Kit in a 96-well plate format.

### 6.8 GEP activity assay

Protease activity was assessed using the chromogenic substrate Z-FLE-pNA (Bachem, 60 µM) in a 200 µL reaction volume containing 10 mM Tris-HCl (pH 7.4). Absorbance at 405 nm was measured every minute for 30 minutes at 37°C with continuous orbital shaking using a Tecan Spark microplate reader (Tecan).

For supernatant-based activity assays, overnight cultures of isolates were diluted 1:100 in LB and grown at 37°C and 180 rpm. At an OD_600_ of 0.8, cultures were centrifuged at 5,000 × g for 20 minutes and filtered through a 0.2 µm sterile filter. Protease activity was assessed by adding 50 µL of filtered supernatant to the reaction described above.

For recombinant protein-based activity assays, proteins were first activated by incubation with thermolysin (Sigma-Aldrich, final concentration 5 µg/mL) in 10 mM Tris-HCl (pH 7.4) supplemented with 2 mM CaCl_2_ for one hour at 37°C with shaking (25), reaching a final recombinant protein concentration of 50 µg/mL in a 100 µL total volume. Thermolysin was subsequently inactivated by the addition of EDTA to a final concentration of 5 mM. Controls without recombinant protein and without thermolysin were included for each protease tested. Protease activity was then assessed by adding activated recombinant protein (2.5 µg/mL final concentration) to the reaction described above.

### 6.9 Proteomics

#### 6.9.1 Peptide clean-up

Recombinant PAR1 N-terminus (166 µg/mL) was incubated with activated recombinant protein (10 µg/mL) in 10 mM Na_2_HPO_4_ for 30 minutes at 37°C in a total volume of 75 µL. The peptide mixtures were acidified by adding 10% trifluoroacetic acid (TFA) to a final concentration of 0.5% and desalted on C18 StageTips based on the protocol of Rappsilber et al., 2007 (62) with some modifications. Three C18 Empore disk punches were packed into 200 µL pipette tips. All applied solutions were passed through the tip by centrifugation. Tips were prewetted with 100 µL methanol, followed by 100 µL 80% acetonitrile (ACN), 0,1% TFA, and equilibrated with 100 µL 0.1% TFA by spinning at 376 × g for 4 - 6 min. Acidified peptide solutions were applied, and spun at 271 × g till passed through. Bound peptides were washed with 100 µL 0.1% TFA, and eluted with 2 x 30 µL 40% ACN, 0.1% TFA into PCR tubes. Eluates were reduced to dryness in a vacuum centrifuge and taken up in 20 µL 2% ACN, 0.1% TFA.

#### 6.9.2 Liquid chromatography-mass spectrometry analysis

LC-MS analysis was performed on a Vanquish Neo UHPLC system (Thermo Scientific) coupled to an Orbitrap Exploris 480 mass spectrometer (Thermo Scientific). The system was equipped with a Nanospray Flex ion source (Thermo Scientific), and a Column Heater (IonOpticks) connected to a Heater Controller (IonOpticks).

Peptides were loaded onto a trap column (PepMap Neo C18 5 mm × 300 µm, 5 μm particle size, Thermo Scientific) using 0.1% TFA as mobile phase, and separated on an analytical column (Aurora Ultimate XT C18, 25 cm × 75 µm, 1.7 µm particle size, IonOpticks), applying a linear gradient starting with a mobile phase of 98% solvent A (0.1% formic acid and 2% solvent B (80% acetonitrile, 0.08% formic acid), increasing to 35% solvent B over 30 min at a flow rate of 300 nL/min. The analytical column was heated to 50°C. The mass spectrometer was operated in data-dependent acquisition (DDA) mode, with 1.2 s MS1 cycle time. Survey scans were acquired from 375-1500 m/z with lock mass enabled, normalized AGC target of 100%, resolution of 60,000. The most intense precursor ions (charge states +2 to +6) were selected for fragmentation using an isolation window of 1.4 m/z. Selected ions were analyzed with a maximum fill time of 100 ms, normalized AGC target of 100%, and resolution of 30,000 after HCD fragmentation with normalized collision energy of 30%. Monoisotopic precursor selection (MIPS) was set to “peptide” mode, the intensity threshold to 2.5E4, and selected precursors were dynamically excluded for 10 seconds with isotope exclusion enabled.

#### 6.9.3 Mass spectrometry data analysis

MS raw data were analyzed with FragPipe (23.1), using MSFragger (4.3) (63), IonQuant (1.11.11) (64), and Philosopher (5.1.2) (65). The default FragPipe workflow for label free quantification was used, except “Normalize intensity across runs” and “Match between runs” was turned off. Cleavage specificity was set to GluC/P, with 12 missed cleavages allowed. Maximal peptides size was set to 80AA and 8000 Da. The protein FDR was set to 1%. Oxidation of methionine and N-terminal protein acetylation were specified as variable modifications, with a maximum of 3 variable modifications allowed per peptide. Cysteine was set to “not modified”. MS2 spectra were searched against the *E*. *coli* 1 protein per gene reference proteome from Uniprot (Proteome ID: UP000000625, release 2025.01), concatenated with a database of common laboratory contaminants (release 2025_01, https://github.com/maxperutzlabs-ms/perutz-ms-contaminants), and entries for the substrate and enzymes. Computational analysis was performed using Python and the Python library MsReport (version 0.0.31; source code: https://github.com/hollenstein/msreport) (66).

### 6.10 Monitoring of epithelial barrier function with the xCELLigence system

xCELLigence experiments were performed as previously described (68). Briefly, primary nasal epithelial cells obtained from four healthy donors (Ethics Committee approval no. EK Nr. 2240/2021) were cultured in collagen type 1 (Corning) coated flasks containing bronchial epithelial growth medium (BEGM; BulletKit medium; Lonza Group Ltd., Basel, Switzerland) at 37°C in a humidified atmosphere with 5% CO₂ until confluence was reached (7-10 days). Subsequently, cells were seeded into collagen/fibronectin-coated wells of E-Plate 16 plates of the xCELLigence Real-Time Cell Analysis (RTCA) dual-purpose (DP) system (ACEA Biosciences, San Diego, USA) at a density of 1 × 10⁵ cells/mL. After cells reached a cell index of 8-12, test substances were added: V8 (Sigma-Aldrich) at 10 and 2.5 µg/ml and expressed V8, Esp, SplB, Csp, and Hsp with or without Thermolysin treatment at a final concentration of 10 µg/mL. The Buffer control was treated with Thermolysin. Impedance-based measurements were recorded every 30 minutes for up to 200 hours. Cell index values were normalized as previously described.

## 7 Acknowledgements

This research was funded in whole or in part by the Austrian Science Fund (FWF) [10.55776/P36339] and FWF cluster of excellence (CoE7) “Microbiomes Drive Planetary Health” [10.55776/CoE7]. For the purpose of open access, the author has applied a CC BY public copyright license to any Author Accepted Manuscript version arising from this submission.

## 10 Supplementary Figures

**Figure S1.**
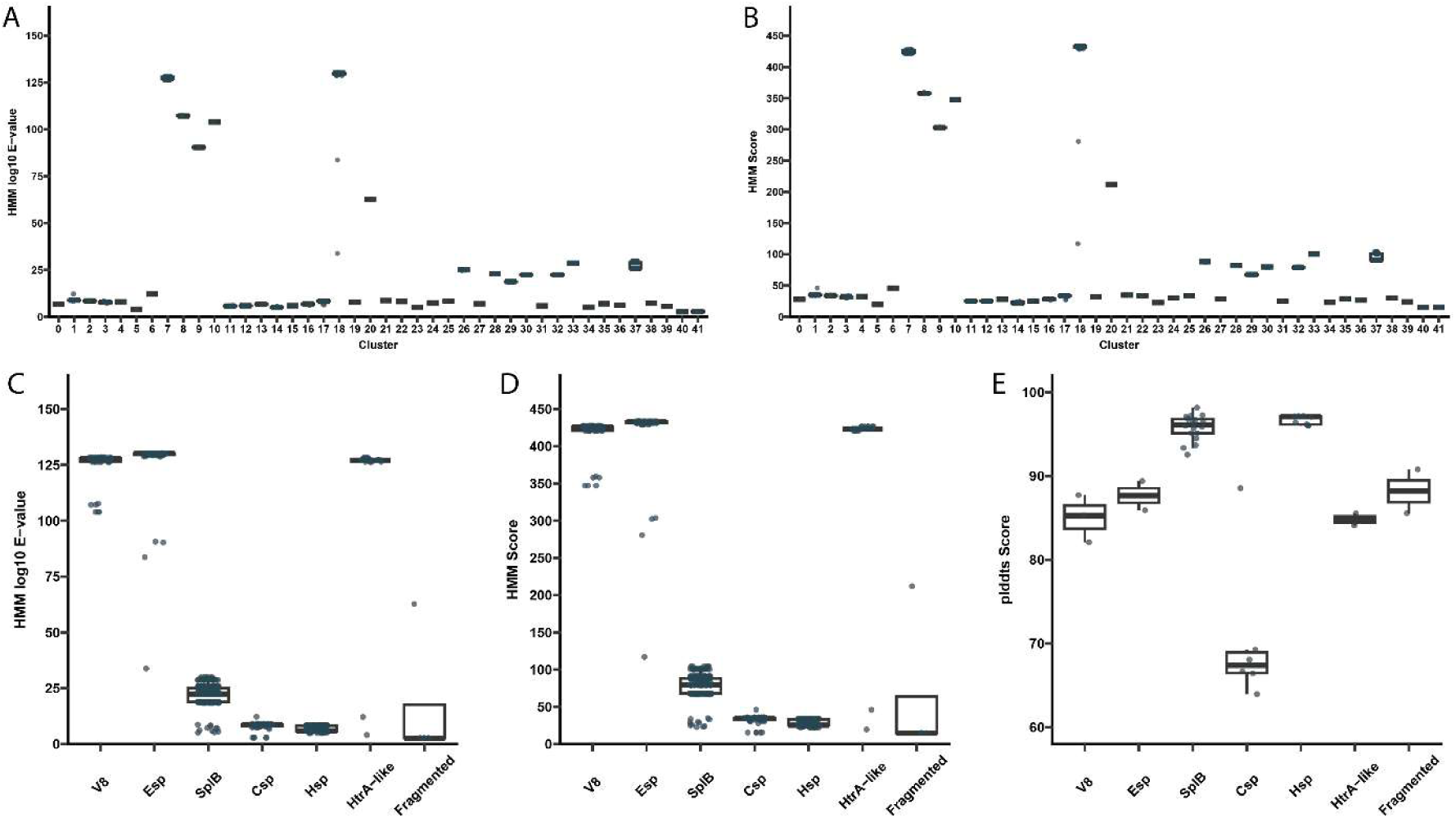
Quality metrics of glutamyl endopeptidase homologs identified by HMM-based screening. (a-b) Identified clusters (CD-HIT) and their (a) distribution of HMM e-values (log₁₀-transformed) and (b) HMM scores. (c-d) Clusters grouped by active site classifications (AlphaFold2-based) and their (c) distribution of HMM e-values (log₁₀-transformed) and (d) HMM scores. (e) pLDDT confidence scores by active site classifications (AlphaFold2-based).

**Figure S2.**
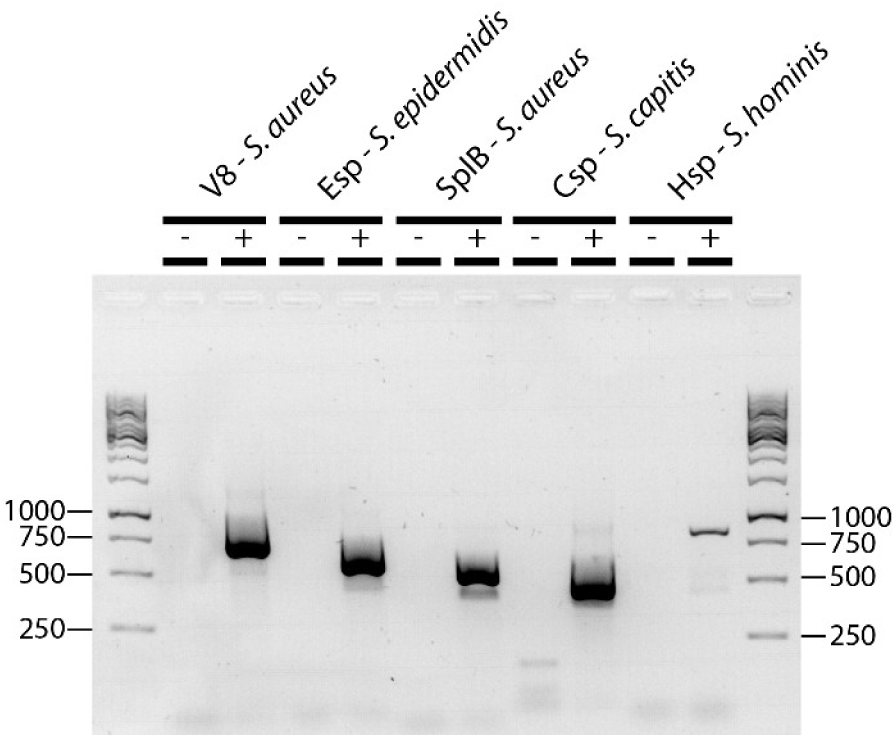
Gene presence confirmation. Confirmation of gene presence in candidate GEP homolog isolates by PCR with their respective size in base pairs confirmed through Sanger sequencing. (-) indicates a PCR reaction without DNA/RNA. (+) indicates a PCR reaction with DNA.

**Figure S3.**
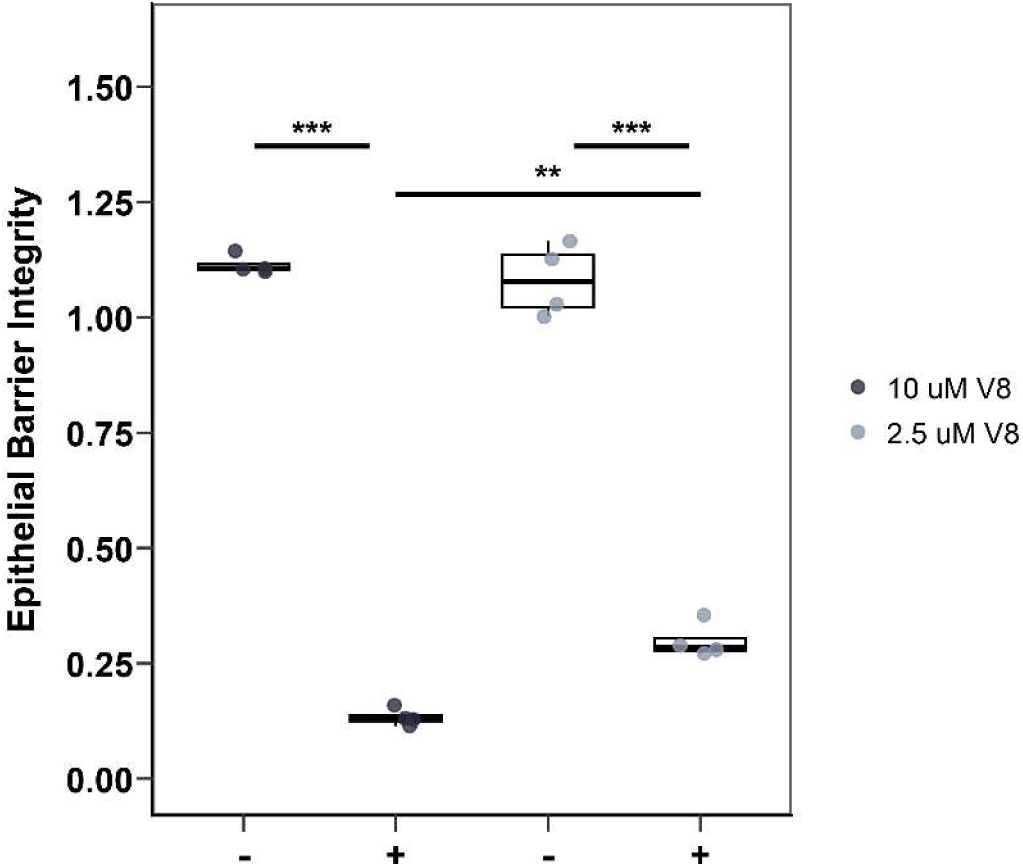
Staphylococcal proteases impair epithelial barrier integrity. Box plots showing the cell index at the 48-hour time point for cells treated with buffer (indicated as -) or a commercially available V8 protease (indicated as +) at 10 µM or 2.5 µM. Differences were assessed by two-way ANOVA with Tukey’s *post-hoc* test. Significance levels: * *p* < 0.05, ** *p* < 0.01, *** *p* < 0.001. n = 2 technical replicates, n = 2 patients.

## Supplementary Table

**Table S1.**
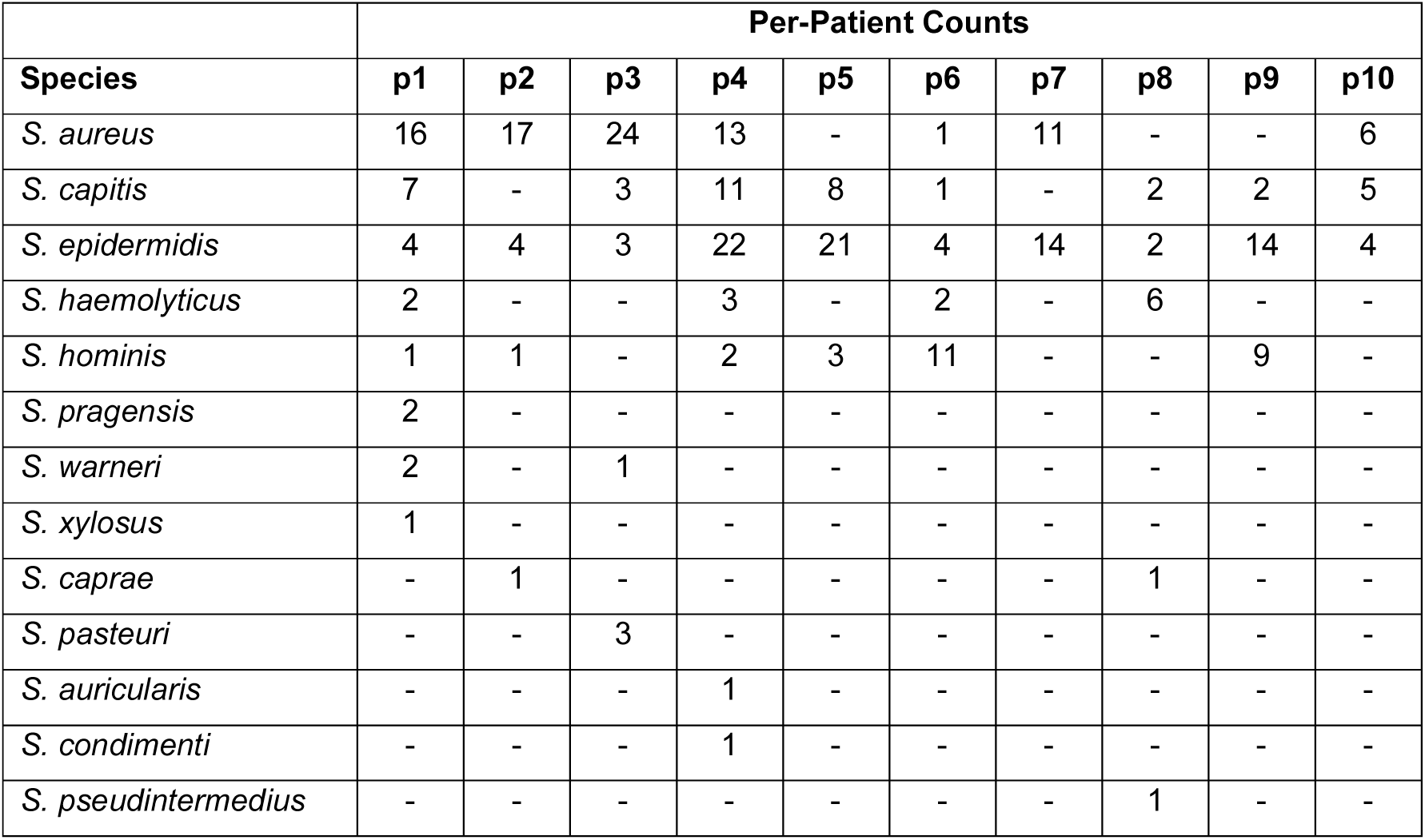
Staphylococcal species isolated from the skin of atopic dermatitis patients. Overview of the 13 *Staphylococcus* species identified across 10 atopic dermatitis patients (p1–p10). Values indicate the number of isolates recovered per species and patient. Species not detected in a patient are indicated by “-“.

**Table S2.**
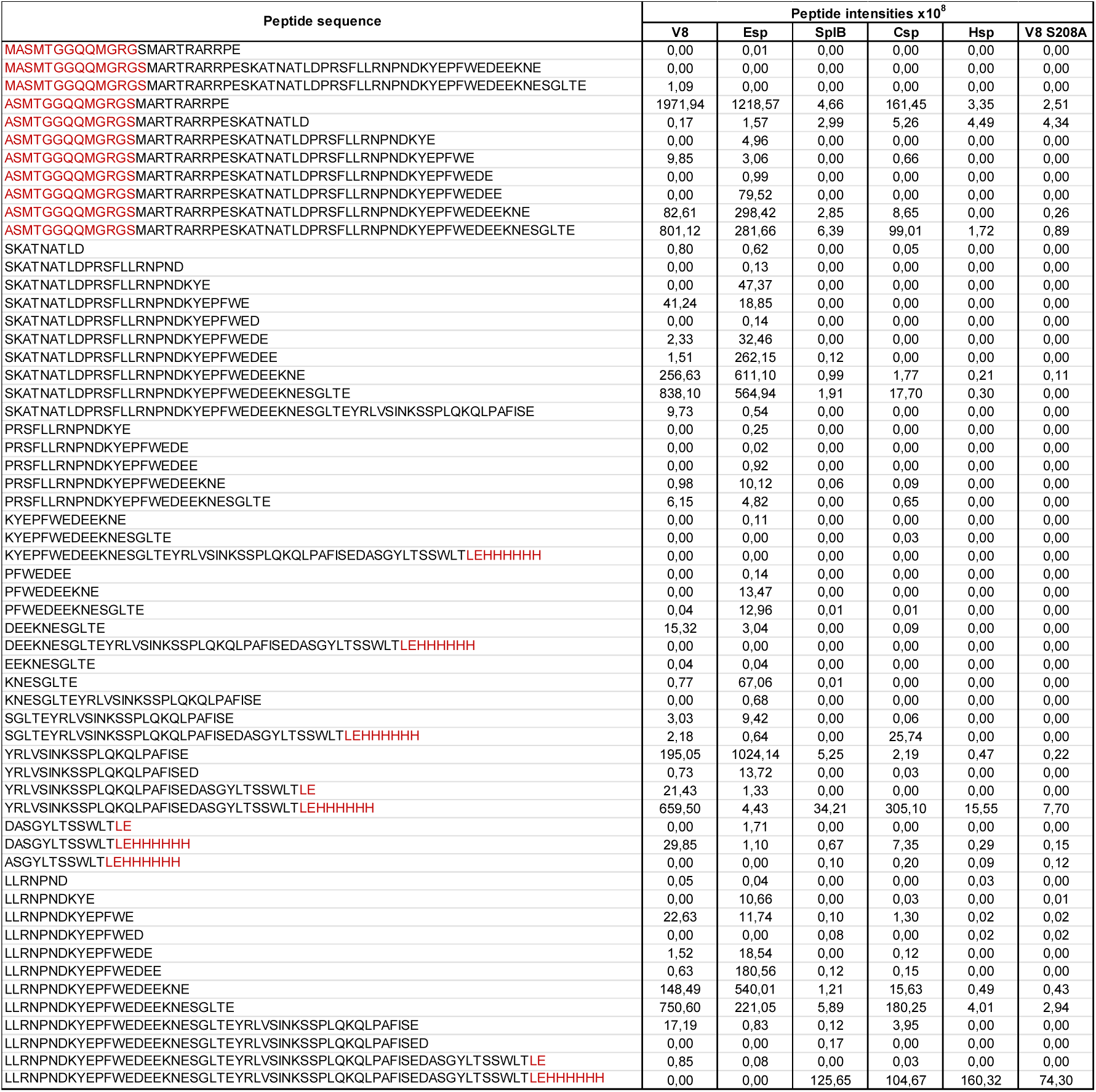
Peptide fragments and their intensity for each protease. Peptide sequences and their intensity after incubation of PAR1 tethered ligand with the respective protease. Amino acids in red result from the expressed His6-tagged construct and are not native to

**Table S3.** BLAST-based genomic screening of V8, Esp, and Csp across publicly available *Staphylococcus* genomes. Results of BLAST searches using V8, Esp, and Csp as reference protein sequences against publicly available *Staphylococcus* genome sequences. Only species yielding at least one hit are shown. For each species-gene combination, the number of detected hits is reported alongside mean sequence identity, query coverage, and log E-value (mean ± SD). NA values for SD indicate that only a single sequence was detected for that species-gene combination.

| Species (n) | Gene | Hits | Identity<br>(Mean ± SD) | Coverage<br>(Mean ± SD) | Log E-Value<br>(Mean ± SD) |
| --- | --- | --- | --- | --- | --- |
| *S. aureus* (9659) | Esp | 1 | 99.64 ± NA | 100 ± NA | 323.31 ± NA |
| *S. aureus* (9659) | SplB | 7037 | 96.63 ± 9.28 | 99.98 ± 0.67 | 169.16 ± 17.44 |
| *S. aureus* (9659) | V8 | 9548 | 96.84 ± 1.43 | 98.82 ± 4.15 | 323.27 ± 2.31 |
| *S. auricularis* (9) | Hsp | 9 | 49.51 ± 0.86 | 84.56 ± 4.67 | 80.21 ± 1.63 |
| *S. capitis* (113) | Hsp | 111 | 69.01 ± 0.23 | 100 ± 0 | 143.77 ± 0.49 |
| *S. capitis* (113) | Csp | 106 | 99.67 ± 0.6 | 100 ± 0 | 172.54 ± 0.93 |
| *S. caprae* (62) | Esp | 61 | 61.6 ± 2.76 | 88.39 ± 9.5 | 106.34 ± 4.71 |
| *S. caprae* (62) | Csp | 61 | 92.74 ± 1.05 | 100 ± 0 | 159.65 ± 2.1 |
| *S. condimentii* (7) | Hsp | 7 | 53.31 ± 0 | 99 ± 0 | 99.72 ± 0.01 |
| *S. condimentii* (7) | Csp | 7 | 76.32 ± 0 | 99 ± 0 | 134.65 ± 0.07 |
| *S. epidermidis* (700) | Esp | 688 | 99.75 ± 0.53 | 99.95 ± 0.83 | 323.09 ± 5.75 |
| *S. epidermidis* (700) | Hsp | 532 | 50.11 ± 0.25 | 98.93 ± 1.16 | 91.47 ± 1.09 |
| *S. haemolyticus* (308) | Hsp | 301 | 73.36 ± 0.25 | 99.95 ± 0.62 | 156.08 ± 1.33 |
| *S. haemolyticus* (308) | Csp | 304 | 89.81 ± 0.28 | 100 ± 0 | 159.72 ± 0.77 |
| *S. hominis* (136) | Hsp | 133 | 99.35 ± 0.24 | 100 ± 0 | 323.31 ± 0 |
| *S. pasteurii* (34) | V8 | 34 | 65.49 ± 1.31 | 83.94 ± 0.34 | 138.66 ± 1.39 |
| *S. pasteurii* (34) | Hsp | 32 | 66.63 ± 1.17 | 91.66 ± 2.07 | 123.87 ± 0.52 |
| *S. pragensis* (1) | Hsp | 1 | 82.41 ± NA | 100 ± NA | 323.31 ± NA |
| *S. pseudintermedius* (263) | SplB | 10 | 45.38 ± 0 | 99 ± 0 | 69.87 ± 0.01 |
| *S. warneri* (76) | V8 | 75 | 73.61 ± 0.31 | 82 ± 0 | 147.37 ± 0.55 |
| *S. warneri* (76) | Hsp | 65 | 63 ± 0.21 | 98 ± 0 | 126.63 ± 1.69 |

**Table S4.**
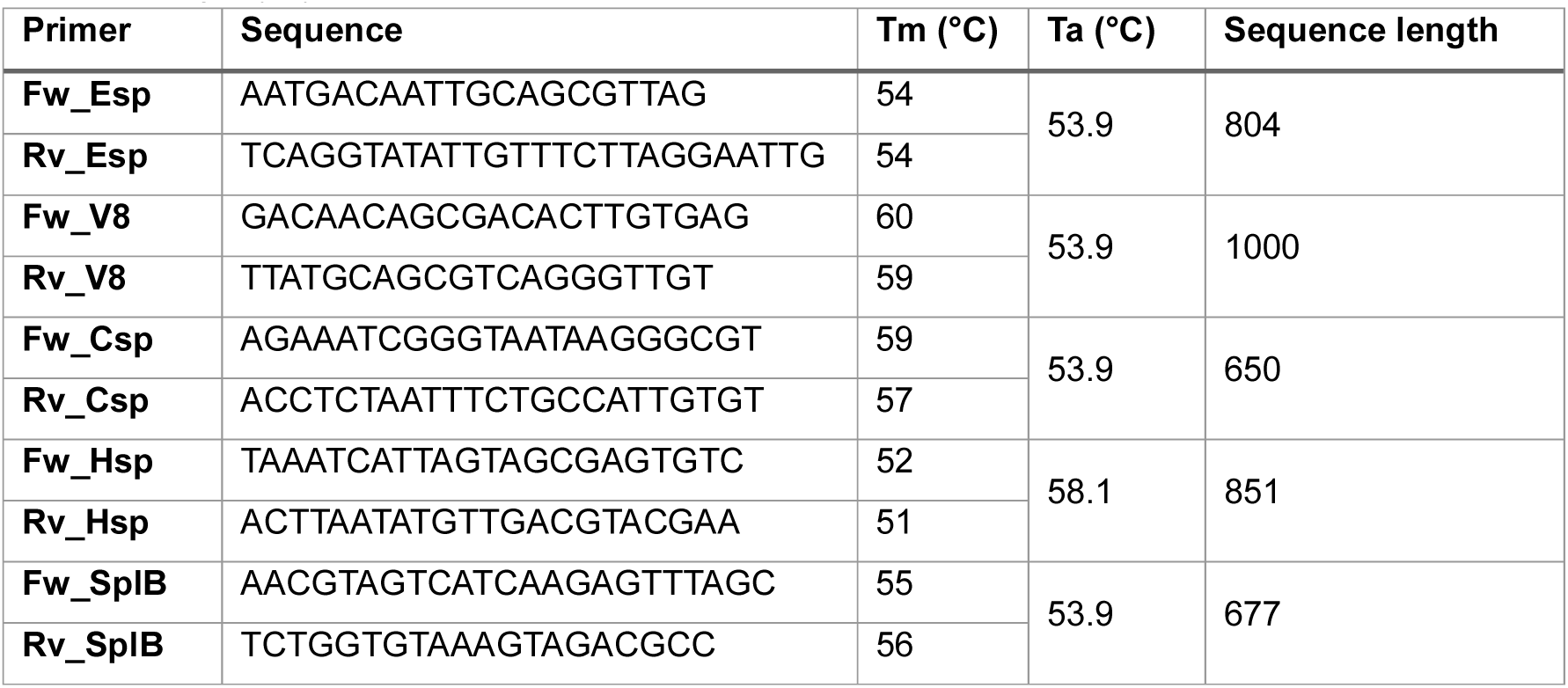
Primers used for reverse transcriptase PCR (RT-PCR) analysis. Table with forward (Fw) and reverse (Rv) primer pairs designed for RT-PCR amplification of five target genes including the nucleotide sequence, melting temperature (Tm, °C), use annealing temperature and amplicon length (bp).

## References

1. Bylund S, Kobyletzki LB, Svalstedt M, Svensson Å. 2020. Prevalence and Incidence of Atopic Dermatitis: A Systematic Review. Acta Derm Venereol 100:adv00160. doi:10.2340/00015555-3510.

2. Weidinger S, Novak N. 2016. Atopic dermatitis. The Lancet 387:1109–1122. doi:10.1016/S0140-6736(15)00149-X.

3. Weidinger S, Beck LA, Bieber T, Kabashima K, Irvine AD. 2018. Atopic dermatitis. Nat Rev Dis Primers 4:1. doi:10.1038/s41572-018-0001-z.

4. Czarnowicki T, He H, Krueger JG, Guttman-Yassky E. 2019. Atopic dermatitis endotypes and implications for targeted therapeutics. Journal of Allergy and Clinical Immunology 143:1–11. doi:10.1016/j.jaci.2018.10.032.

5. Chng KR, Tay ASL, Li C, Ng AHQ, Wang J, Suri BK, Matta SA, McGovern N, Janela B, Wong, Xuan Fei Colin C., Sio YY, Au BV, Wilm A, Sessions PF de, Lim TC, Tang MBY, Ginhoux F, Connolly JE, Lane EB, Chew FT, Common JEA, Nagarajan N. 2016. Whole metagenome profiling reveals skin microbiome-dependent susceptibility to atopic dermatitis flare. Nat Microbiol 1:16106. doi:10.1038/nmicrobiol.2016.106.

6. Gonzalez ME, Schaffer JV, Orlow SJ, Gao Z, Li H, Alekseyenko AV, Blaser MJ. 2016. Cutaneous microbiome effects of fluticasone propionate cream and adjunctive bleach baths in childhood atopic dermatitis. J Am Acad Dermatol 75:481–493.e8. doi:10.1016/j.jaad.2016.04.066.

7. Kong HH, Oh J, Deming C, Conlan S, Grice EA, Beatson MA, Nomicos E, Polley EC, Komarow HD, Murray PR, Turner ML, Segre JA. 2012. Temporal shifts in the skin microbiome associated with disease flares and treatment in children with atopic dermatitis. Genome Res 22:850–859. doi:10.1101/gr.131029.111.

8. Totté JEE, van der Feltz WT, Hennekam M, van Belkum A, van Zuuren EJ, Pasmans SGMA. 2016. Prevalence and odds of Staphylococcus aureus carriage in atopic dermatitis: a systematic review and meta-analysis. Br J Dermatol 175:687–695. doi:10.1111/bjd.14566.

9. Grice EA, Kong HH, Conlan S, Deming CB, Davis J, Young AC, Bouffard GG, Blakesley RW, Murray PR, Green ED, Turner ML, Segre JA. 2009. Topographical and temporal diversity of the human skin microbiome. Science 324:1190–1192. doi:10.1126/science.1171700.

10. Tomassi A de, Reiter A, Reiger M, Rauer L, Rohayem R, CK-CARE Study Group, Traidl-Hoffmann C, Neumann AU, Hülpüsch C. 2023. Combining 16S Sequencing and qPCR Quantification Reveals Staphylococcus aureus Driven Bacterial Overgrowth in the Skin of Severe Atopic Dermatitis Patients. Biomolecules 13:1030. doi:10.3390/biom13071030.

11. Deng L, Costa F, Blake KJ, Choi S, Chandrabalan A, Yousuf MS, Shiers S, Dubreuil D, Vega-Mendoza D, Rolland C, Deraison C, Voisin T, Bagood MD, Wesemann L, Frey AM, Palumbo JS, Wainger BJ, Gallo RL, Leyva-Castillo J-M, Vergnolle N, Price TJ, Ramachandran R, Horswill AR, Chiu IM. 2023. S. aureus drives itch and scratch-induced skin damage through a V8 protease-PAR1 axis. Cell 186:5375–5393.e25. doi:10.1016/j.cell.2023.10.019.

12. Akiyama H, Toi Y, Kanzaki H, Tada J, Arata J. 1996. Prevalence of producers of enterotoxins and toxic shock syndrome toxin-1 among Staphylococcus aureus strains isolated from atopic dermatitis lesions. Arch Dermatol Res 288:418–420. doi:10.1007/BF02507115.

13. Bunikowski R, Mielke ME, Skarabis H, Worm M, Anagnostopoulos I, Kolde G, Wahn U, Renz H. 2000. Evidence for a disease-promoting effect of Staphylococcus aureus-derived exotoxins in atopic dermatitis. Journal of Allergy and Clinical Immunology 105:814–819. doi:10.1067/mai.2000.105528.

14. Mcfadden JP, Noble WC, Camp RD. 1993. Superantigenic exotoxin-secreting potential of staphylococci isolated from atopic eczematous skin. Br J Dermatol 128:631–632. doi:10.1111/j.1365-2133.1993.tb00257.x.

15. Severn MM, Williams MR, Shahbandi A, Bunch ZL, Lyon LM, Nguyen A, Zaramela LS, Todd DA, Zengler K, Cech NB, Gallo RL, Horswill AR. 2022. The Ubiquitous Human Skin Commensal Staphylococcus hominis Protects against Opportunistic Pathogens. mBio 13:e0093022. doi:10.1128/mbio.00930-22.

16. Byrd AL, Belkaid Y, Segre JA. 2018. The human skin microbiome. Nat Rev Microbiol 16:143–155. doi:10.1038/nrmicro.2017.157.

17. Cau L, Williams MR, Butcher AM, Nakatsuji T, Kavanaugh JS, Cheng JY, Shafiq F, Higbee K, Hata TR, Horswill AR, Gallo RL. 2021. Staphylococcus epidermidis protease EcpA can be a deleterious component of the skin microbiome in atopic dermatitis. Journal of Allergy and Clinical Immunology 147:955–966.e16. doi:10.1016/j.jaci.2020.06.024.

18. Severn MM, Horswill AR. 2023. Staphylococcus epidermidis and its dual lifestyle in skin health and infection. Nat Rev Microbiol 21:97–111. doi:10.1038/s41579-022-00780-3.

19. Williams MR, Costa SK, Zaramela LS, Khalil S, Todd DA, Winter HL, Sanford JA, O’Neill AM, Liggins MC, Nakatsuji T, Cech NB, Cheung AL, Zengler K, Horswill AR, Gallo RL. 2019. Quorum sensing between bacterial species on the skin protects against epidermal injury in atopic dermatitis. Sci Transl Med 11. doi:10.1126/scitranslmed.aat8329.

20. Wang B, McHugh BJ, Qureshi A, Campopiano DJ, Clarke DJ, Fitzgerald JR, Dorin JR, Weller R, Davidson DJ. 2017. IL-1β-Induced Protection of Keratinocytes against Staphylococcus aureus-Secreted Proteases Is Mediated by Human β-Defensin 2. J Invest Dermatol 137:95–105. doi:10.1016/j.jid.2016.08.025.

21. Yokoi K-J, Kuzuwa S, Iwasaki S-I, Yamakawa A, Taketo A, Kodaira K-I. 2016. Aureolysin of Staphylococcus warneri M accelerates its proteolytic cascade, and participates in biofilm formation. Bioscience, biotechnology, and biochemistry 80:1238–1242. doi:10.1080/09168451.2016.1148576.

22. Calkins S, Couger MB, Jackson C, Zandler J, Hudgins GC, Hanafy RA, Budd C, French DP, Hoff WD, Youssef N. 2016. Draft genome sequence of Staphylococcus hominis strain Hudgins isolated from human skin implicates metabolic versatility and several virulence determinants. Genomics Data 10:91–96. doi:10.1016/j.gdata.2016.10.003.

23. Simpson EL, Villarreal M, Jepson B, Rafaels N, David G, Hanifin J, Taylor P, Boguniewicz M, Yoshida T, Benedetto A de, Barnes KC, Leung DYM, Beck LA. 2018. Patients with Atopic Dermatitis Colonized with Staphylococcus aureus Have a Distinct Phenotype and Endotype. J Invest Dermatol 138:2224–2233. doi:10.1016/j.jid.2018.03.1517.

24. Polgár L. 20XX-. Chapter 560 - Catalytic Mechanisms of Serine and Threonine Peptidases, p. 2524–2534. In Rawlings ND (ed), Handbook of proteolytic enzymes, 3rd ed. Academic Press, Amsterdam, Heidelberg.

25. Nemoto TK, Ohara-Nemoto Y, Ono T, Kobayakawa T, Shimoyama Y, Kimura S, Takagi T. 2008. Characterization of the glutamyl endopeptidase from Staphylococcus aureus expressed in Escherichia coli. The FEBS Journal 275:573–587. doi:10.1111/j.1742-4658.2007.06224.x.

26. Jenul C, Horswill AR. 2019. Regulation of Staphylococcus aureus Virulence. Microbiol Spectr 7. doi:10.1128/microbiolspec.GPP3-0031-2018.

27. Gallo RL, Horswill AR. 2024. Staphylococcus aureus: The Bug Behind the Itch in Atopic Dermatitis. J Invest Dermatol 144:950–953. doi:10.1016/j.jid.2024.01.001.

28. Hansen KK, Saifeddine M, Hollenberg MD. 2004. Tethered ligand-derived peptides of proteinase-activated receptor 3 (PAR3) activate PAR1 and PAR2 in Jurkat T cells. Immunology 112:183–190. doi:10.1111/j.1365-2567.2004.01870.x.

29. Benedetto A de, Rafaels NM, McGirt LY, Ivanov AI, Georas SN, Cheadle C, Berger AE, Zhang K, Vidyasagar S, Yoshida T, Boguniewicz M, Hata T, Schneider LC, Hanifin JM, Gallo RL, Novak N, Weidinger S, Beaty TH, Leung DYM, Barnes KC, Beck LA. 2011. Tight junction defects in patients with atopic dermatitis. Journal of Allergy and Clinical Immunology 127:773–86.e1-7. doi:10.1016/j.jaci.2010.10.018.

30. Nakatsuji T, Chen TH, Two AM, Chun KA, Narala S, Geha RS, Hata TR, Gallo RL. 2016. Staphylococcus aureus Exploits Epidermal Barrier Defects in Atopic Dermatitis to Trigger Cytokine Expression. J Invest Dermatol 136:2192–2200. doi:10.1016/j.jid.2016.05.127.

31. Murphy J, Ramezanpour M, Stach N, Dubin G, Psaltis AJ, Wormald P-J, Vreugde S. 2018. Staphylococcus Aureus V8 protease disrupts the integrity of the airway epithelial barrier and impairs IL-6 production in vitro. The Laryngoscope 128:E8–E15. doi:10.1002/lary.26949.

32. Azghani AO, Gray LD, Johnson AR. 1993. A bacterial protease perturbs the paracellular barrier function of transporting epithelial monolayers in culture. Infect Immun 61:2681–2686. doi:10.1128/iai.61.6.2681-2686.1993.

33. Nakamura Y, Takahashi H, Takaya A, Inoue Y, Katayama Y, Kusuya Y, Shoji T, Takada S, Nakagawa S, Oguma R, Saito N, Ozawa N, Nakano T, Yamaide F, Dissanayake E, Suzuki S, Villaruz A, Varadarajan S, Matsumoto M, Kobayashi T, Kono M, Sato Y, Akiyama M, Otto M, Matsue H, Núñez G, Shimojo N. 2020. Staphylococcus Agr virulence is critical for epidermal colonization and associates with atopic dermatitis development. Sci Transl Med 12. doi:10.1126/scitranslmed.aay4068.

34. Abdurrahman G, Pospich R, Steil L, Gesell Salazar M, Izquierdo González JJ, Normann N, Mrochen D, Scharf C, Völker U, Werfel T, Bröker BM, Roesner LM, Gómez-Gascón L. 2024. The extracellular serine protease from Staphylococcus epidermidis elicits a type 2-biased immune response in atopic dermatitis patients. Front Immunol 15:1352704. doi:10.3389/fimmu.2024.1352704.

35. Olson ME, Todd DA, Schaeffer CR, Paharik AE, van Dyke MJ, Büttner H, Dunman PM, Rohde H, Cech NB, Fey PD, Horswill AR. 2014. Staphylococcus epidermidis agr quorum-sensing system: signal identification, cross talk, and importance in colonization. J Bacteriol 196:3482–3493. doi:10.1128/JB.01882-14.

36. Byrd AL, Deming C, Cassidy SKB, Harrison OJ, Ng W-I, Conlan S, Belkaid Y, Segre JA, Kong HH. 2017. Staphylococcus aureus and Staphylococcus epidermidis strain diversity underlying pediatric atopic dermatitis. Sci Transl Med 9. doi:10.1126/scitranslmed.aal4651.

37. Le KY, Park MD, Otto M. 2018. Immune Evasion Mechanisms of Staphylococcus epidermidis Biofilm Infection. Front. Microbiol. 9:359. doi:10.3389/fmicb.2018.00359.

38. Uçkay I, Pittet D, Vaudaux P, Sax H, Lew D, Waldvogel F. 2009. Foreign body infections due to Staphylococcus epidermidis. Annals of medicine 41:109–119. doi:10.1080/07853890802337045.

39. Perez-Riverol Y, Bandla C, Kundu DJ, Kamatchinathan S, Bai J, Hewapathirana S, John NS, Prakash A, Walzer M, Wang S, Vizcaíno JA. 2025. The PRIDE database at 20 years: 2025 update. Nucleic acids research 53:D543–D553. doi:10.1093/nar/gkae1011.

40. Quast C, Pruesse E, Yilmaz P, Gerken J, Schweer T, Yarza P, Peplies J, Glöckner FO. 2013. The SILVA ribosomal RNA gene database project: improved data processing and web-based tools. Nucleic acids research 41:D590–6. doi:10.1093/nar/gks1219.

41. Yilmaz P, Parfrey LW, Yarza P, Gerken J, Pruesse E, Quast C, Schweer T, Peplies J, Ludwig W, Glöckner FO. 2014. The SILVA and “All-species Living Tree Project (LTP)” taxonomic frameworks. Nucleic acids research 42:D643–8. doi:10.1093/nar/gkt1209.

42. Kolmogorov M, Yuan J, Lin Y, Pevzner PA. 2019. Assembly of long, error-prone reads using repeat graphs. Nat Biotechnol 37:540–546. doi:10.1038/s41587-019-0072-8.

43. Gurevich A, Saveliev V, Vyahhi N, Tesler G. 2013. QUAST: quality assessment tool for genome assemblies. Bioinformatics 29:1072–1075. doi:10.1093/bioinformatics/btt086.

44. Mikheenko A, Prjibelski A, Saveliev V, Antipov D, Gurevich A. 2018. Versatile genome assembly evaluation with QUAST-LG. Bioinformatics 34:i142–i150. doi:10.1093/bioinformatics/bty266.

45. Parks DH, Imelfort M, Skennerton CT, Hugenholtz P, Tyson GW. 2015. CheckM: assessing the quality of microbial genomes recovered from isolates, single cells, and metagenomes. Genome Res 25:1043–1055. doi:10.1101/gr.186072.114.

46. Chaumeil P-A, Mussig AJ, Hugenholtz P, Parks DH. 2019. GTDB-Tk: a toolkit to classify genomes with the Genome Taxonomy Database. Bioinformatics 36:1925–1927. doi:10.1093/bioinformatics/btz848.

47. Chaumeil P-A, Mussig AJ, Hugenholtz P, Parks DH. 2022. GTDB-Tk v2: memory friendly classification with the genome taxonomy database. Bioinformatics 38:5315–5316. doi:10.1093/bioinformatics/btac672.

48. Seemann T. 2014. Prokka: rapid prokaryotic genome annotation. Bioinformatics 30:2068–2069. doi:10.1093/bioinformatics/btu153.

49. Eddy SR. 2011. Accelerated Profile HMM Searches. PLOS Computational Biology 7:e1002195. doi:10.1371/journal.pcbi.1002195.

50. Sievers F, Wilm A, Dineen D, Gibson TJ, Karplus K, Li W, Lopez R, McWilliam H, Remmert M, Söding J, Thompson JD, Higgins DG. 2011. Fast, scalable generation of high-quality protein multiple sequence alignments using Clustal Omega. Mol Syst Biol 7:539. doi:10.1038/msb.2011.75.

51. Li W, Godzik A. 2006. Cd-hit: a fast program for clustering and comparing large sets of protein or nucleotide sequences. Bioinformatics 22:1658–1659. doi:10.1093/bioinformatics/btl158.

52. Fu L, Niu B, Zhu Z, Wu S, Li W. 2012. CD-HIT: accelerated for clustering the next-generation sequencing data. Bioinformatics 28:3150–3152. doi:10.1093/bioinformatics/bts565.

53. Teufel F, Almagro Armenteros JJ, Johansen AR, Gíslason MH, Pihl SI, Tsirigos KD, Winther O, Brunak S, Heijne G von, Nielsen H. 2022. SignalP 6.0 predicts all five types of signal peptides using protein language models. Nat Biotechnol 40:1023–1025. doi:10.1038/s41587-021-01156-3.

54. Jumper J, Evans R, Pritzel A, Green T, Figurnov M, Ronneberger O, Tunyasuvunakool K, Bates R, Žídek A, Potapenko A, Bridgland A, Meyer C, Kohl SAA, Ballard AJ, Cowie A, Romera-Paredes B, Nikolov S, Jain R, Adler J, Back T, Petersen S, Reiman D, Clancy E, Zielinski M, Steinegger M, Pacholska M, Berghammer T, Bodenstein S, Silver D, Vinyals O, Senior AW, Kavukcuoglu K, Kohli P, Hassabis D. 2021. Highly accurate protein structure prediction with AlphaFold. Nature 596:583–589. doi:10.1038/s41586-021-03819-2.

55. Jones P, Binns D, Chang H-Y, Fraser M, Li W, McAnulla C, McWilliam H, Maslen J, Mitchell A, Nuka G, Pesseat S, Quinn AF, Sangrador-Vegas A, Scheremetjew M, Yong S-Y, Lopez R, Hunter S. 2014. InterProScan 5: genome-scale protein function classification. Bioinformatics 30:1236–1240. doi:10.1093/bioinformatics/btu031.

56. Camacho C, Coulouris G, Avagyan V, Ma N, Papadopoulos J, Bealer K, Madden TL. 2009. BLAST+: architecture and applications. BMC Bioinformatics 10:421. doi:10.1186/1471-2105-10-421.

57. 2026. R: The R Project for Statistical Computing. https://www.r-project.org/. Accessed 3 June, 2026.

58. Wickham H, Averick M, Bryan J, Chang W, McGowan L, François R, Grolemund G, Hayes A, Henry L, Hester J, Kuhn M, Pedersen T, Miller E, Bache S, Müller K, Ooms J, Robinson D, Seidel D, Spinu V, Takahashi K, Vaughan D, Wilke C, Woo K, Yutani H. 2019. Welcome to the Tidyverse. JOSS 4:1686. doi:10.21105/joss.01686.

59. H. Pagès, P. Aboyoun, R. Gentleman, and S. DebRoy. 2017. Biostrings. Bioconductor.

60. Champely S. 2006. CRAN: Contributed Packages.

61. Signorell A. 2014. CRAN: Contributed Packages.

62. Rappsilber J, Mann M, Ishihama Y. 2007. Protocol for micro-purification, enrichment, pre-fractionation and storage of peptides for proteomics using StageTips. Nat Protoc 2:1896–1906. doi:10.1038/nprot.2007.261.

63. Kong AT, Leprevost FV, Avtonomov DM, Mellacheruvu D, Nesvizhskii AI. 2017. MSFragger: ultrafast and comprehensive peptide identification in mass spectrometry-based proteomics. Nat Methods 14:513–520. doi:10.1038/nmeth.4256.

64. Yu F, Haynes SE, Nesvizhskii AI. 2021. IonQuant Enables Accurate and Sensitive Label-Free Quantification With FDR-Controlled Match-Between-Runs. Molecular & Cellular Proteomics 20:100077. doi:10.1016/j.mcpro.2021.100077.

65. Da Veiga Leprevost F, Haynes SE, Avtonomov DM, Chang H-Y, Shanmugam AK, Mellacheruvu D, Kong AT, Nesvizhskii AI. 2020. Philosopher: a versatile toolkit for shotgun proteomics data analysis. Nat Methods 17:869–870. doi:10.1038/s41592-020-0912-y.

66. David Hollenstein. 2025. hollenstein/msreport: MsReport v0.0.31 - Transformer and Normalizer update. Zenodo.

67. Hollenstein DM, Hartl M. 2025. hollenstein/xlsxreport: v0.1.0. Zenodo.

68. Waltl EE, Selb R, Eckl-Dorna J, Mueller CA, Cabauatan CR, Eiwegger T, Resch-Marat Y, Niespodziana K, Vrtala S, Valenta R, Niederberger V. 2018. Betamethasone prevents human rhinovirus- and cigarette smoke- induced loss of respiratory epithelial barrier function. Sci Rep 8:9688. doi:10.1038/s41598-018-27022-y.

